# Projected expansion in climatic suitability for *Trypanosoma cruzi*, the etiological agent of Chagas Disease, and five widespread *Triatoma* species by 2070

**DOI:** 10.1101/490508

**Authors:** Matthew D. Nichols, Chris J. Butler, Wayne D. Lord, Michelle L. Haynie

## Abstract

The vector-borne parasite *Trypanosoma cruzi* infects seven million individuals globally and causes chronic cardiomyopathy and gastrointestinal diseases. Recently, *T. cruzi* has emerged in the southern United States. It is crucial for disease surveillance efforts to detail regions that present favorable climatic conditions for *T. cruzi* and vector establishment. We used MaxEnt to develop an ecological niche model for *T. cruzi* and five widespread *Triatoma* vectors based on 546 published localities within the United States. We modeled regions of current potential *T. cruzi* and *Triatoma* distribution and then regions projected to have suitable climatic conditions by 2070. Regions with suitable climatic conditions for the study organisms are predicted to increase within the United States. Our findings agree with the hypothesis that climate change will facilitate the expansion of tropical diseases throughout temperate regions and suggest climate change will influence the expansion of *T. cruzi* and *Triatoma* vectors in the United States.

**Author summary:** *Trypanosoma cruzi* is a vector-borne parasite and the etiological agent of Chagas disease, which is an emerging infectious disease in the United States. Current trends suggest climate change can influence the distribution of vector-borne diseases through increases in suitable climatic conditions for vectors and pathogens. This influence can lead to tropical disease emergence in temperate regions. Currently, the best method for estimating distributional changes and potential disease emergence in both well studied and underrepresented areas is ecological niche modeling. We developed an ecological niche model to better understand the potential future *T. cruzi* and *Triatoma* distribution in the United States and in Oklahoma, which is an underrepresented state. Our ecological niche model incorporated 546 published localities and 19 bioclimatic variables. Our model predicts a potential range expansion of *T. cruzi* and widespread *Triatoma* vectors throughout Oklahoma and the United States. Our model presents a better understanding of the future epidemiology of *T. cruzi* and *Triatoma* vectors in Oklahoma and the United States and contributes to disease surveillance efforts by predicting areas of potential range expansion.

## Introduction

*Trypanosoma cruzi* is a vector-borne hemoflagellate parasite and the etiological agent of American Trypanosomiasis, also known as Chagas disease. Currently, *T. cruzi* infects seven million people across 43 countries and up to 40% will develop Chagas disease, which causes significant cardiomyopathy in adults and children, tissue fibrosis, lethargy, gastrointestinal diseases such as megaesophagus and megacolon, and 10,000 deaths annually [1–5]. *Trypanosoma cruzi* is transmitted when infected hematophagous triatomines (Hemiptera: Reduviidae: Triatominae) feed on a host and defecate onto the host skin or mucous membranes, thereby allowing the parasite to enter the host via the bite wound [4]. *Trypanosoma cruzi* is known to infect over 400 mammalian species and is prevalent within wildlife populations in endemic regions where triatomine vectors occur [6–9].

Current trends suggest global climate change will result in an expansion of tropical diseases, notably vector-borne diseases, throughout temperate regions [10–11]. Examples of concern include schistosomiasis, onchocerciasis, dengue fever, lymphatic filariasis, African and American trypanosomiases, yellow fever, and other mosquito and tick-transmitted diseases of humans [10–11]. By 2050, the climate of England will again be suitable for endemic malaria [12]. As climatic temperature and humidity increase, conditions can influence disease morbidity, most notably the length of transmission [13]. For example, malaria transmission can increase to epidemic, hypoendemic, mesoendemic, hyperendemic, and holoendemic levels [11], [13]. Furthermore, floods and droughts caused by climate change can instigate disease outbreaks by creating breeding grounds for insects whose desiccated eggs remain viable and hatch in still water [10].

Over the last 100 years, global mean surface air temperatures over land and oceans have increased beyond any period in the past 40 million years [14]. One report modeled species extinction and estimated between 33% and 58% of all species will become extinct by 2050 under scenarios of maximum expected climate change [15]. Additionally, projections estimate climate change will influence the distribution and expansion of tropical diseases, notably vector-borne diseases, throughout temperate regions [10–11]. Because of the effects of observed changes in the distribution and phenology of organisms caused by warming in the 20th century, it is important to model how climate change may influence infectious disease ecology within domestic, wildlife, and human populations [16–17].

Ecological niche modeling (ENM) is a valuable tool for understanding the geographic ecology of a species. ENM estimates the dimensions of species’ ecological niches, which is the space within which a species can maintain populations with immigration [18–19]. ENM predicts the fundamental and realized niches of species by relating point occurrence data of species to environmental factors [20–21]. Through machine learning, a customized genetic algorithm predicts and confirms the following: high predictive ability of the approach regarding species’ distributions, the ability to predict species’ potential distributions across scenarios of change on ecologic and evolutionary time scales, the ability to predict the course of species’ invasions, the capacity to understand and predict the geographic outcomes of species’ interactions, and useful insight into various other aspects of species’ distributional ecology [18–30].

ENM is an essential tool for understanding the geographic dimensions of the risk of transmission of *T. cruzi*. ENM facilitates the exploration of geographic and ecologic phenomena based on known occurrences of the study species [17], [31–32]. ENM is used to better understand the epidemiology of *T. cruzi* through niche characterization of triatomines, and relationships between vector and reservoir distributions [19], [33]. Studies of the geographic distribution of *T. cruzi* and widespread *Triatoma* vectors are crucial for understanding the epidemiologic aspects of *T. cruzi* transmission and must be taken into consideration when focusing control efforts and disease surveillance in underrepresented areas [32].

Maximum Entropy (MaxEnt) modeling uses species presence-only data and environmental conditions to estimate the distribution of a species [33]. The basis of MaxEnt is to minimize the relative entropy between two probability densities defined in covariate space [33]. By predicting the entire geographic range in which a species might occur, the realized niche does not limit the fundamental niche. This approach can be used to assess the relative importance of specific environmental factors to a species distribution, locate areas of current suitable habitat, and project changes in its distribution over time [33].

Recently, increasing reports of autochthonous vectorial transmission suggest *T. cruzi* is endemic in the United States and enzootic transmission cycles are more prevalent than expected [5], [34–43]. Currently, there are 29 states with reports of *T. cruzi* and triatomine vectors [5]. There are four reports of *T. cruzi* in Oklahoma wildlife, but its endemicity within the state is underrepresented when compared to other southern states [34–36], [43].

The World Health Organization instigated a vector control and eradication program in Latin America which has substantially decreased transmission in rural regions of Latin American and reduced disease incidence by 94% in the Southern Cone countries [44]. Decades of successful vector eradication campaigns in Latin America, along with regional programs focused on reducing vector infestation within human dwellings and blood screening, decreased the total prevalence of Chagas disease from >16 million to 8 million people [4], [45–46].

Since 2007, *T. cruzi* control efforts in Latin America were unified to combat the globalization of Chagas disease, which addresses the immigration of infected individuals into non-endemic countries and the potential for non-vectorial transmission routes [3], [5]. Despite the successful eradication programs and the effort to minimize globalization, *T. cruzi* infections have spread globally through human immigration [5], [45], [47]. The United States has an estimated 300,000 infected individuals, most of which are immigrants from areas endemic to *T. cruzi* [5]. With the emergence of *T. cruzi* in the United States; the widespread distribution of vectors within the United States; and projected effects of climate change on parasite, vector, and reservoir distribution; there is a need to determine the potential current and future distributions of these organisms to better understand their influence on the epidemiology of *T. cruzi* in the United States.

## Methods

We used Maxent and the ‘ENMeval’ package in R [48] to model the current and projected distribution of *T. cruzi* as well as five widespread potential vectors: *Triatoma gerstaeckeri, T. indictiva, T. lecticularia, T. protracta*, and *T. sanguisuga* [49–51]. We collected documented occurrences of these six species from published records and incorporated records that met one or more of the following criteria: 1) documentation of the parasite in accepted endemic areas; 2) multiple cases of human infection (three or more) within an area; and 3) reports of the parasite found in intermediate or definitive hosts (S1 and S2 Tables). We included 546 published location data points of *T. cruzi* and the five *Triatoma* species in this study (S1 and S2 Tables) and downloaded elevation and 19 bioclimatic variables from WorldClim (Table 1; [52–53]; http://www.worldclim.org/) at a resolution of 10 arc minutes (400 km^2^). We avoided model overfitting using a regularization approach which introduced a penalty for an increase in model complexity [50], [54], and the small sample corrected variant of Akaike’s information criterion (AICc) scores was used to evaluate the regularization of models [55]. Future climate conditions for 2070 using the IPCC 5 data from WorldClim [52] were used to project the potential future distribution of the six species of interest at 10 arc minutes using the model that best predicted the current distribution of each species. Projected distributions were computed based on the IPCC scenario RCP 8.5 (emissions increase throughout the 21st century) using the ACCESS1.3 general circulation models.

**Table 1.** Summary of bioclimatic variables used in this study.

| Variable | Definition |
| --- | --- |
| BIO 1 | Annual mean temperature |
| BIO 2 | Mean diurnal range (mean of monthly [max temp – min temp]) |
| BIO 3 | Isothermality (BIO 2 / BIO 7) x 100 |
| BIO 4 | Temperature seasonality (standard deviation x 100) |
| BIO 5 | Max temperature of warmest month |
| BIO 6 | Min temperature of coldest month |
| BIO 7 | Temperature annual range (BIO 5 – BIO 6) |
| BIO 8 | Mean temperature of wettest quarter |
| BIO 9 | Mean temperature of driest quarter |
| BIO 10 | Mean temperature of warmest quarter |
| BIO 11 | Mean temperature of coldest quarter |
| BIO 12 | Annual precipitation |
| BIO 13 | Precipitation of wettest month |
| BIO 14 | Precipitation of driest month |
| BIO 15 | Precipitation seasonality (coefficient of variation) |
| BIO 16 | Precipitation of wettest quarter |
| BIO 17 | Precipitation of driest quarter |
| BIO 18 | Precipitation of warmest quarter |
| BIO 19 | Precipitation of coldest quarter |
| Elevation | Elevation above sea level |

## Results

We obtained location data from 215 published records for *T. cruzi* (S1 Table). Current potential suitable climatic conditions for *T. cruzi* include nearly half of the United States and nearly all of Oklahoma. Areas with suitable climatic conditions for *T. cruzi* are predicted to increase in the United States and Oklahoma by 2070 under the RCP 8.5 scenario (Fig 1; Table 2). We obtained location data from 70 published records for *T. gerstaeckeri* (S2 Table). Areas with suitable climatic conditions for *T. gerstaeckeri* are predicted to increase in the United States by 2070 under the RCP 8.5 scenario (Fig 2; Table 2). At a lower resolution, the potential distribution includes areas of Oklahoma in 2070; however, with a finer resolution, current and future potential distributions do not include areas of Oklahoma (Fig 2). We obtained location data from 12 published records for *T. indictiva* (S2 Table). Areas with suitable climatic conditions for *T. indictiva* are predicted to drastically increase in the central and northern United States by 2070 under the RCP 8.5 scenario (Fig 3). The potential distribution of *T. indictiva* is predicted to increase in Oklahoma by 2070 under the RCP 8.5 scenario (Fig 3; Table 2).

**Table 2.** Mean ± standard deviation projected suitability for the study species. Data generated using MaxEnt.

|  | Current – lower<br>48 | Current –<br>Oklahoma | 2070 RCP 8.5 –<br>lower 48 states | 2070 RCP 8.5 –<br>Oklahoma |
| --- | --- | --- | --- | --- |
| *Trypanosoma*<br>*cruzi* | 0.359 ± 0.389 | 0.995 ± 0.025 | 0.818 ± 0.323 | 1.000 ± 0.000 |
| *Triatoma*<br>*gerstaeckeri* | 0.052 ± 0.135 | 0.110 ± 0.068 | 0.084 ± 0.158 | 0.332 ± 0.162 |
| *Triatoma*<br>*indictiva* | 0.077 ± 0.186 | 0.569 ± 0.240 | 0.249 ± 0.308 | 0.844 ± 0.229 |
| *Triatoma*<br>*lecticularia* | 0.138 ± 0.203 | 0.453 ± 0.142 | 0.333 ± 0.303 | 0.837 ± 0.093 |
| *Triatoma*<br>*protracta* | 0.128 ± 0.218 | 0.161 ± 0.159 | 0.175 ± 0.252 | 0.299 ± 0.255 |
| *Triatoma*<br>*sanguisuga* | 0.263 ± 0.324 | 0.932 ± 0.188 | 0.597 ± 0.405 | 1.000 ± 0.002 |

**Fig 1.**
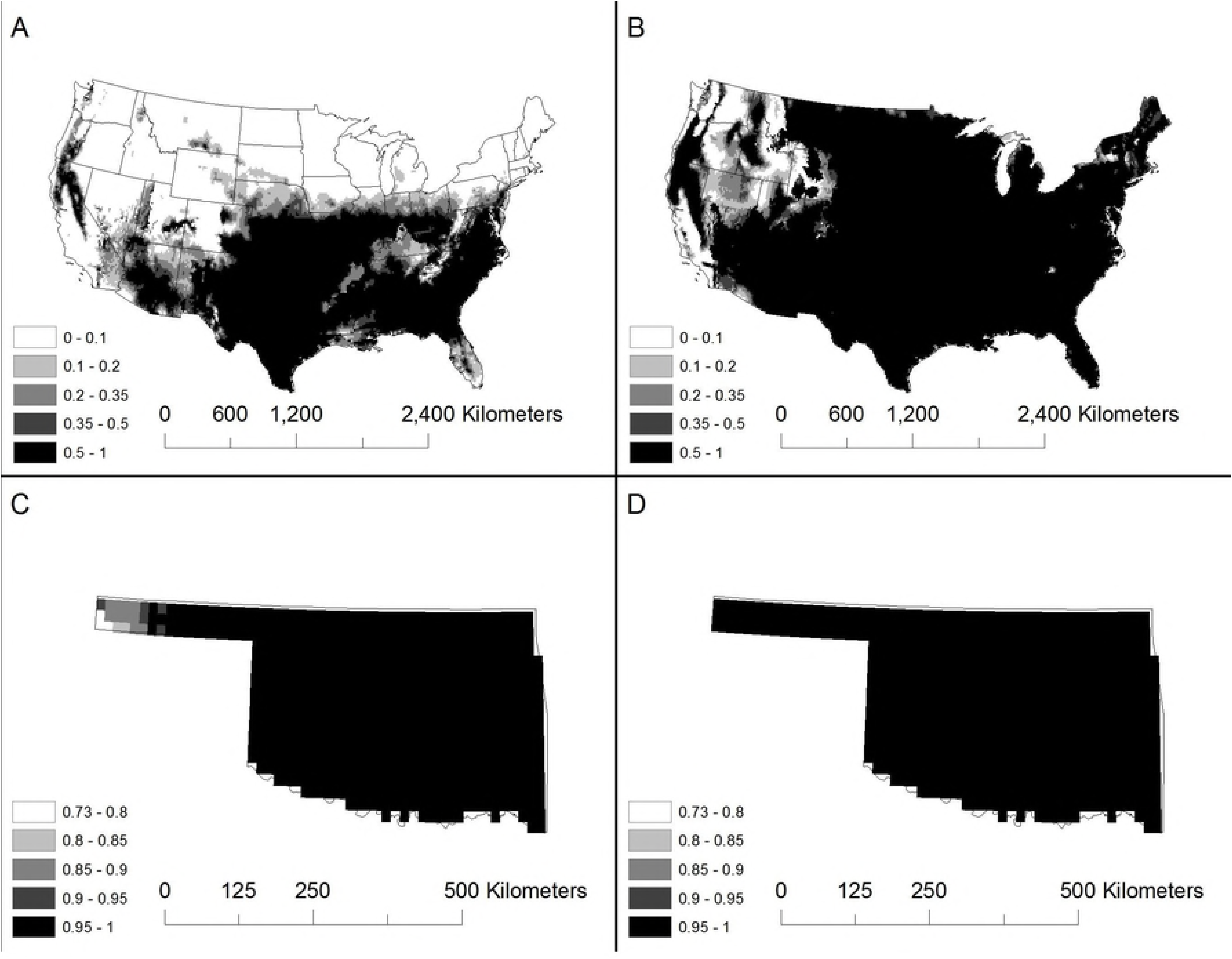
The current potential distribution of *Trypanosoma cruzi* in the United States (A), and the potential distribution by 2070 (B), the current potential distribution in Oklahoma (C), and the potential distribution in OK by 2070 (D). Using a finer resolution for OK, we predict an increase in future potential suitable climatic conditions statewide. Maps were generated using MaxEnt.

**Fig 2.**
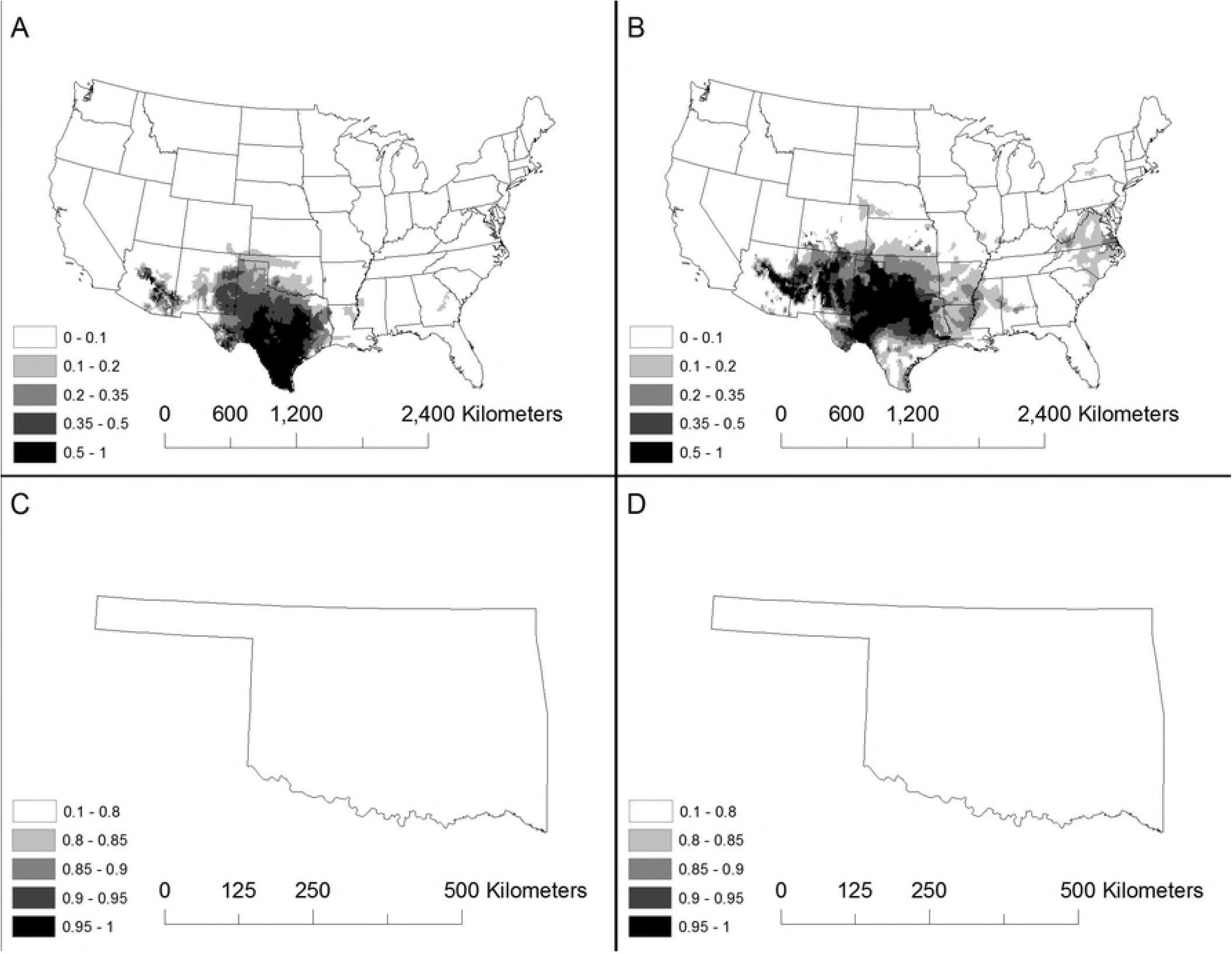
The current potential distribution of *Triatoma gerstaeckeri* in the United States (A), and the potential distribution by 2070 (B), the current potential distribution in Oklahoma (C), and the potential distribution in OK by 2070 (D). Using a finer resolution for OK, we do not predict an increase in future potential suitable climatic conditions by 2070 for this species. We believe this is due to the historic arid environmental conditions preferred by *T. gerstaeckeri*. Maps were generated using MaxEnt.

**Fig 3.**
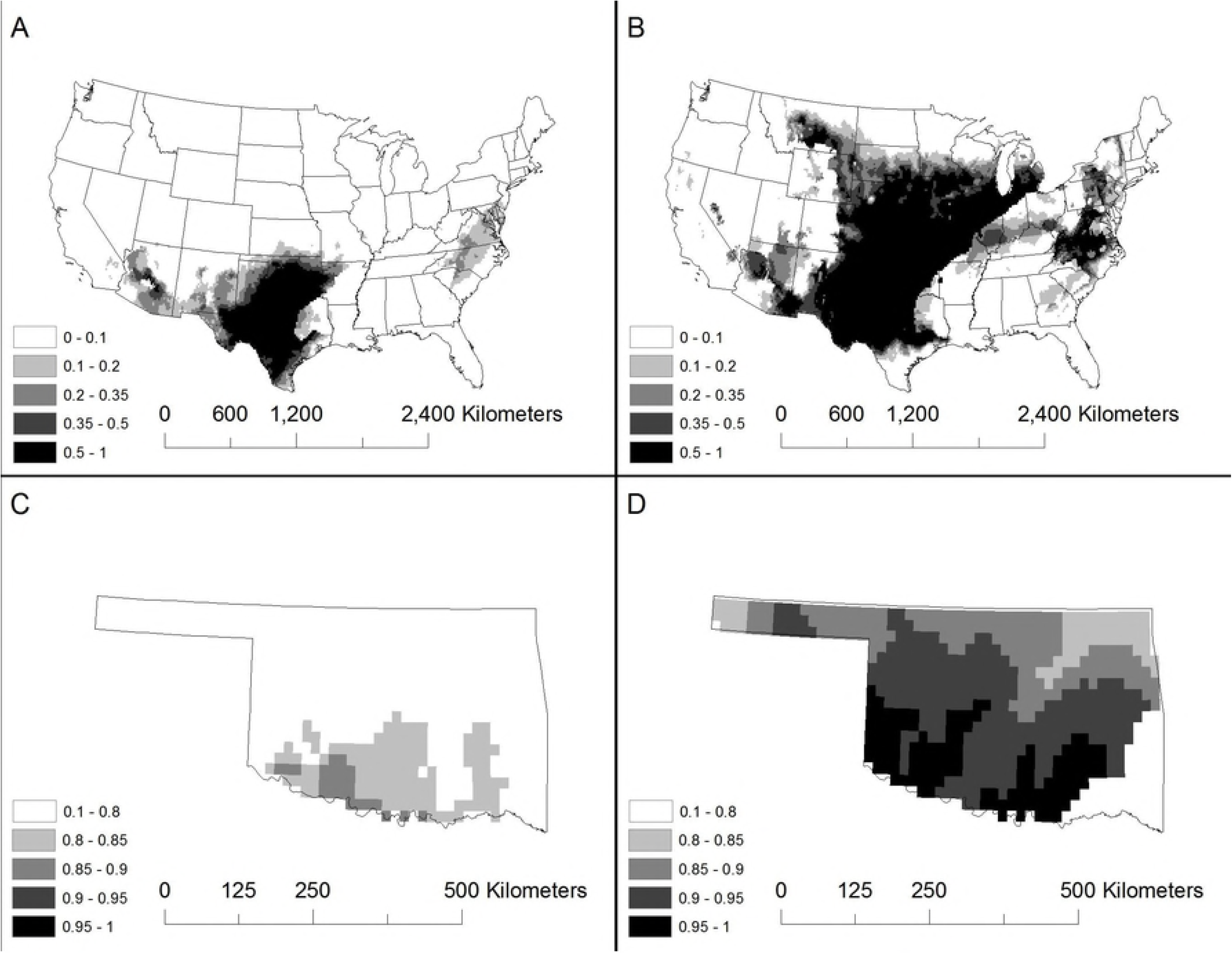
The current potential distribution of *Triatoma indictiva* in the United States (A), and the potential distribution by 2070 (B), the current potential distribution in Oklahoma (C), and the potential distribution in OK by 2070 (D). Using a finer resolution for OK, we predict an increase in future potential suitable climatic conditions for the majority of the state. Maps were generated using MaxEnt.

We obtained location data from 51 published records for *T. lecticularia* (S2 Table). Current areas with potential suitable climatic conditions are found throughout the central and eastern United States (Fig 4). Areas with suitable climatic conditions for *T. lecticularia* are predicted to increase in the United States and Oklahoma by 2070 under the RCP 8.5 scenario (Fig 4; Table 2). We obtained location data from 69 published records for *T. protracta* (S2 Table). Current areas with potential suitable climatic conditions are found throughout the central and western United States (Fig 5). Areas with suitable climatic conditions for *T. protracta* are predicted to increase in the United States and Oklahoma by 2070 under the RCP 8.5 scenario (Fig 5; Table 2). Using a finer resolution for Oklahoma, the potential distribution of *T. protracta* increases into the Oklahoma panhandle by 2070 (Fig 5). We obtained location data from 130 published records for *T. sanguisuga* (S2 Table). Current areas with potential suitable climatic conditions are found throughout the central and eastern United States (Fig 6). Areas with suitable climatic conditions for *T. sanguisuga* are predicted to dramatically increase in the United States and Oklahoma by 2070 under the RCP 8.5 scenario (Fig 6; Table 2). Using a finer resolution for Oklahoma, the potential distribution of *T. sanguisuga* increases statewide by 2070 (Fig 6). The bioclimatic variables that contributed the most to predicting the potential distribution of *T. cruzi* and the *Triatoma* vectors were annual mean temperature, mean diurnal range, annual precipitation, max temperature of the warmest month, and precipitation of the warmest quarter.

**Fig 4.**
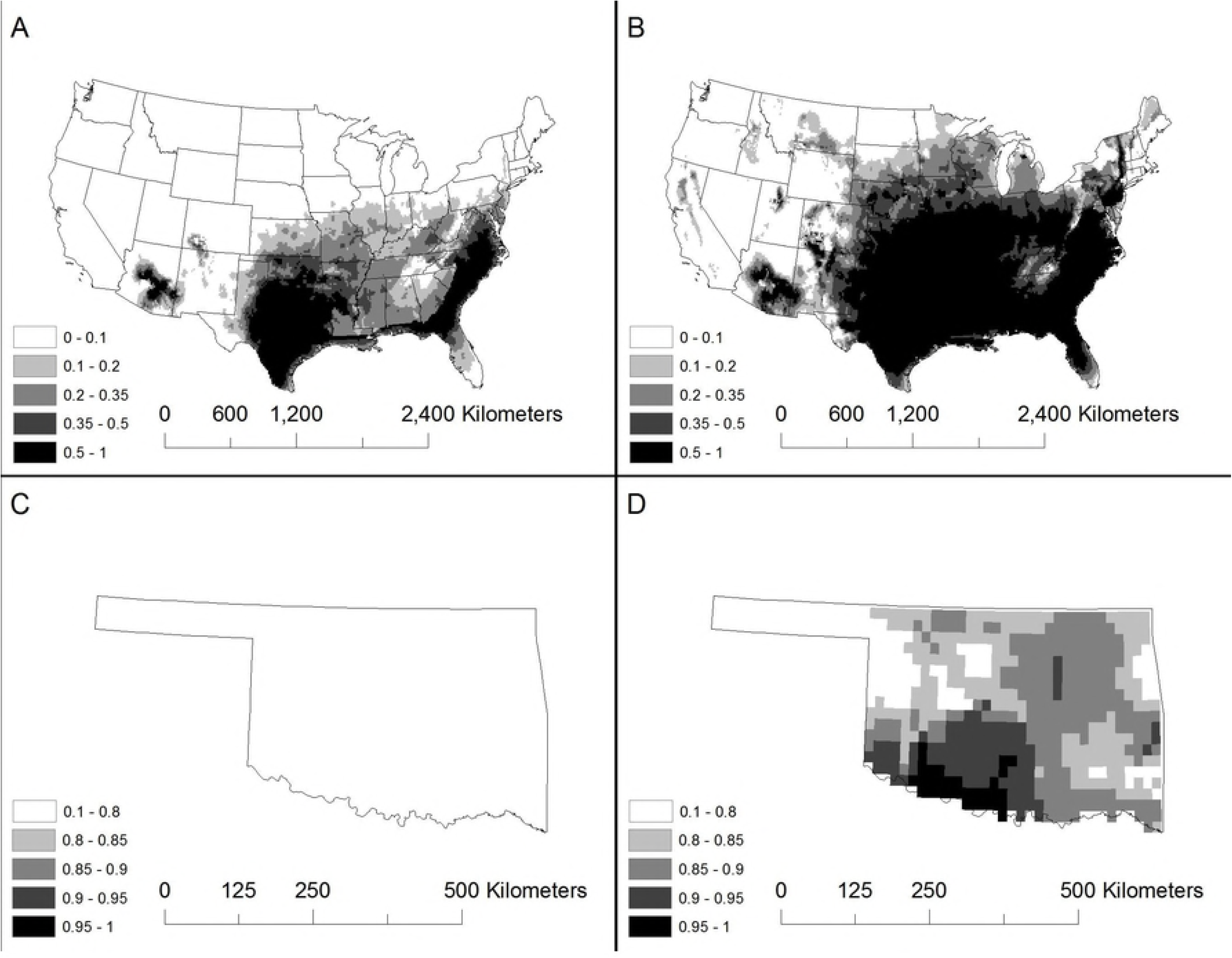
The current potential distribution of *Triatoma lecticularia* in the United States (A), and the potential distribution by 2070 (B), the current potential distribution in Oklahoma (C), and the potential distribution in OK by 2070 (D). Using a finer resolution for OK, we predict an increase in future potential suitable climatic conditions for the majority of the state. Maps were generated using MaxEnt.

**Fig 5.**
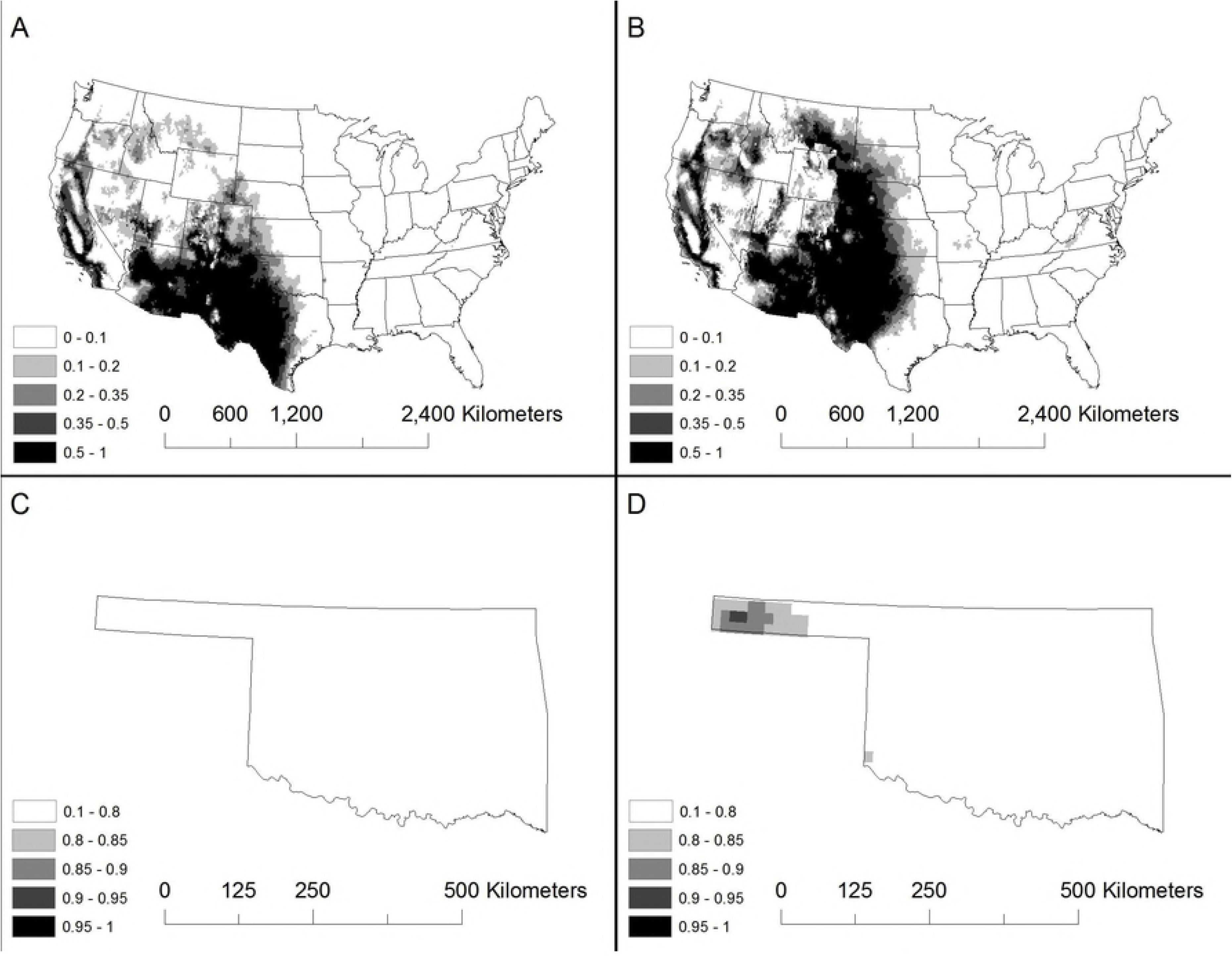
The current potential distribution of *Triatoma protracta* in the United States (A), and the potential distribution by 2070 (B), the current potential distribution in Oklahoma (C), and the potential distribution in OK by 2070 (D). Using a finer resolution for OK, we predict an increase in future potential suitable climatic conditions for the Oklahoma panhandle. Maps were generated using MaxEnt.

**Fig 6.**
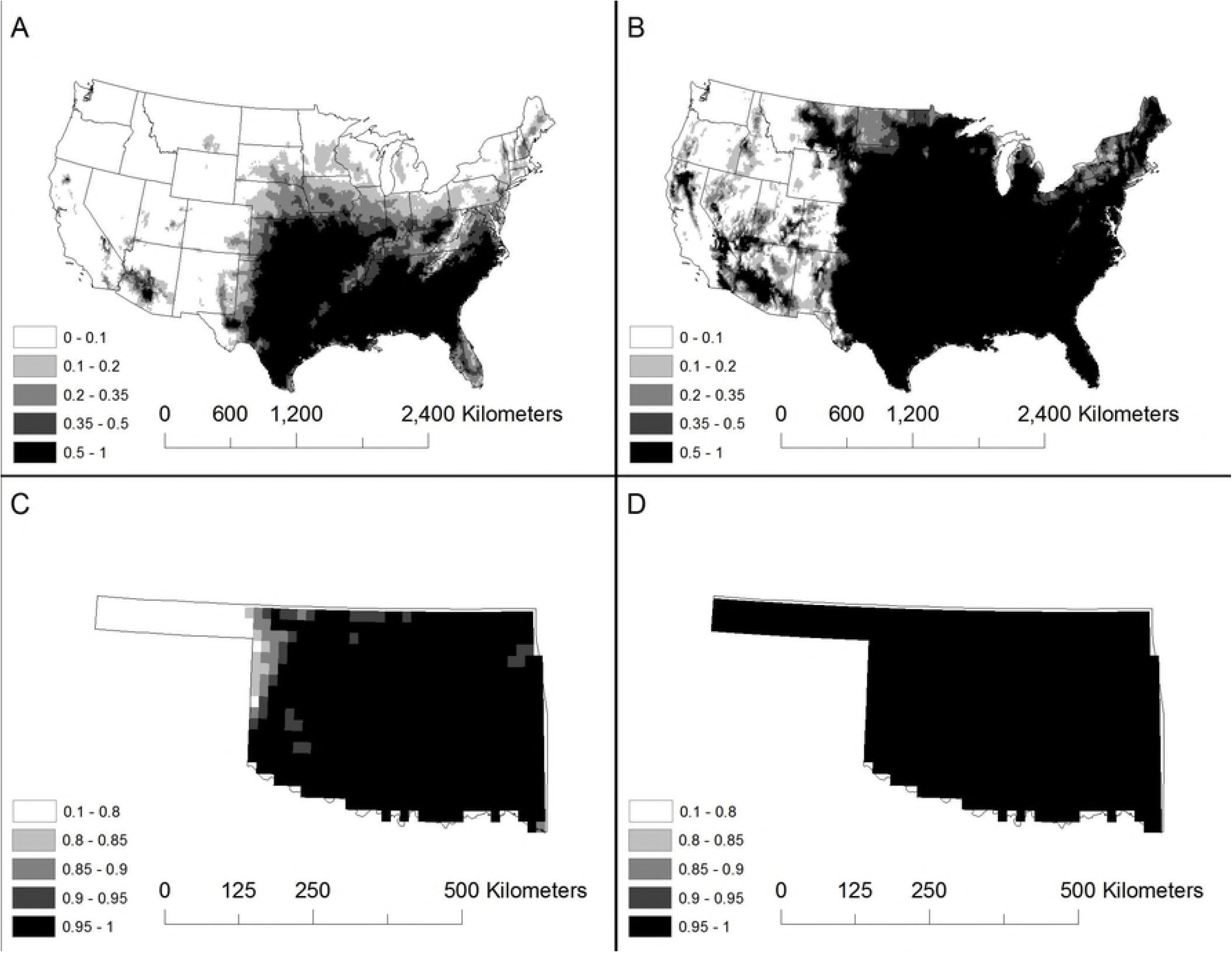
The current potential distribution of *Triatoma sanguisuga* in the United States (A), and the potential distribution by 2070 (B), the current potential distribution in Oklahoma (C), and the potential distribution in OK by 2070 (D). Using a finer resolution for OK, we predict an increase in future potential suitable climatic conditions statewide. Maps were generated using MaxEnt.

## Discussion

Infectious diseases are predicted to emerge in novel foci because of the potential implications of global climate change on the pathogen and vector/host biogeography [10–11]. The influence of climate change will manifest as an increase of disease outbreaks in current regions and expanded transmission risk and disease emergence to novel regions [10], [56–58]. Vector-borne diseases will see marked increases in pathogen and vector distribution. By 2085, total land area favorable for Dengue fever transmission will place up to 60% of the global population at risk for infection [59]. The sandfly, a prominent vector of *Leishmania* parasites, is predicted to expand northward into the United States and can increase transmission risk in novel foci [60]. With the future expansion of pathogens into novel foci, there can be a shift or decline in habitat suitability in current foci, leading to a decline of infections in current foci [11], [17], [61].

Elevating temperature can directly increase the potential for vector-borne diseases and pathogens to increase in disease morbidity [62–64]. For instance, rises in temperature can increase the development time for *Plasmodium, T. cruzi*, and schistosome cercaria [63], [65–67]. Consequently, elevating temperature may affect the pathogen transmission potential and pathogen mortality might increase.

Under the hypothesized IPCC climatic scenarios, our model predicts an overall increase in habitat suitability for *T. cruzi* in the United States, which favors an increase in potential distribution by 2070. The vectors *T. gerstaeckeri, T. indictiva, T. lecticularia, T. protracta,* and *T. sanguisuga* also express this trend, which supports previous literature that vector-borne diseases will spread into temperate regions through increases in suitable vector habitat [10–11]. Additionally, migratory bats, such as the Mexican free-tailed bat (*Tadarida brasiliensis*), may play a unique role in the expansion of *T. cruzi* into temperate regions by migrating between endemic *T. cruzi* regions from Mexico to the Central United States [43].

Historically, *T. cruzi* and *Triatoma* vectors have plagued humans and animal reservoirs in Central and South America, where climate favored parasite transmission and vector biology. Recent molecular evidence suggests that *T. cruzi* evolved from a bat trypanosome in South America approximately 6.5-8.5 million years ago [68–70]. Shortly after trypanosome-infected bats colonized South America 7-10 million years ago, South American humans became infected with *T. cruzi* [71]. The earliest detected human case comes from a 9000-year-old Chinchorro mummy identified via PCR amplification of kinetoplastid DNA sequences [72]. *Trypanosoma cruzi* infected up to 41% of the Chinchorro population located in the Atacama Desert, and this region is where Chagas disease likely originated [72–73]. After the Chinchorro population settled and farmed in regions where sylvatic *T. cruzi* cycles occurred, a domestic transmission cycle emerged [72–75]. The ability of different triatomine vectors, particularly *T. infestans,* to quickly adapt to human dwellings facilitated the domestic *T. cruzi* transmission cycle [76].

Temperature preference can influence the transmission dynamics and epidemiology of *T. cruzi* and competent vectors. Increases in temperature directly increase insect metabolism [77]. In one study, mice were experimentally inoculated with virulent *T. cruzi* strains and subjected to different temperatures [78]. When the mice were kept at 10°C, observed parasitemia became severe after nine days, and all the mice died between the 21st and 26th days [78]. When the mice were kept at 35°C, trypanosomes were undetectable in the blood and from sections of the heart [78]. When the mice were at 26°C, the mice developed a chronic infection [78]. In another study, mice were experimentally infected with a virulent strain, and all mice maintained at 25 ±2°C died 9-15 days post inoculation [65]. These findings suggest a high environmental temperature protected the mice against the virulent effects of *T. cruzi*.

One group of authors examined the influence of temperature on the development of *T. cruzi* while in *Rhodnius prolixus*, which is a common South American vector [67]. The authors hypothesized that the temperature preference of *R. prolixus* also is an optimum temperature range of *T. cruzi*, which is 25.0-25.4°C [79]. At this temperature range, *T. cruzi* has a high *in vitro* growth rate and expresses unrestrained growth which increases the transmission risk. The authors noted a direct relationship between *T. cruzi* parasitemia levels and temperature [67]. When kept at 30°C, *T. cruzi* increased its numbers by 28 times, which doubled its growth rate from 27°C [67]. Lower temperatures affect the endocytic processes of *T. cruzi* epimastigotes and increase *T. cruzi* mortality [80]. At 28°C, *T. cruzi* takes one month to colonize the triatomine intestinal tract, reach the rectum, and differentiate into metacyclic trypomastigotes [81–82]. *Trypanosoma cruzi* infected *R. prolixus* instar molts were delayed by more than 10 days per instar stage [67], [83–84]. Because triatomines only feed after they have molted, it benefits *T. cruzi* to delay their molt and subsequent bloodmeal until *T. cruzi* has colonized, replicated, and is ready to be transmitted, which favors *T. cruzi* transmission potential [67].

Although the potential distribution and habitat suitability may be favorable for disease transmission in the United States, many factors inhibit *T. cruzi* from maintaining a high prevalence within humans in the United States. These include the lack of suitable domestic dwellings for local triatomine vectors to colonize, triatomine expressed zoophilicity, varying and/or delayed triatomine post feeding to defecation time, low virulence of some indigenous *T. cruzi* strains, historic temperate climate, and the possibility of misdiagnosis [83], [85–87].

We present a potential range expansion for *T. cruzi* and five important *Triatoma* species. For this study, we modeled habitat suitability based on 19 bioclimatic variables. Based on our model, we believe the potential significant range expansion for *T. cruzi* and the studied *Triatoma* species, coupled with predicted climate change and low disease surveillance creates a perfect storm for Chagas Disease emergence in the United States.

Future studies should compare the infection rates of important vectors, consider the post-feeding defecation times of each vector, and consider human dwellings and peridomestic animals for precise areas of high-risk transmission potential, which are likely areas of poor housing where vectors can readily colonize. Lastly, we urge physicians, veterinarians, public health officials, and researchers to increase disease surveillance for *T. cruzi* and triatomine vectors to better understand the current and future epidemiology of *T. cruzi,* triatomine vectors, and reservoir hosts in the United States.

## Acknowledgments

We thank Sarah Vrla for her editorial comments. We thank the University of Central Oklahoma College of Mathematics and Sciences for the laboratory space to conduct this research.

## Supporting information

**S1 Table. Reports of *Trypanosoma cruzi* found in intermediate or definitive hosts.**

**S2 Table. Reports of the five *Triatoma* species included in this study.**

**S1 and S2 Table Literature Cited.**

